# Cortical mapping of sensory responses reveals strong brain-state dependence of the late component

**DOI:** 10.1101/2023.10.16.562034

**Authors:** E Montagni, F Resta, N Tort-Colet, A Scaglione, G Mazzamuto, A Destexhe, FS Pavone, AL Allegra Mascaro

## Abstract

Sensory information must be integrated across a distributed brain network for stimulus processing and perception. Recent studies have revealed specific spatiotemporal patterns of cortical activation for the early and late components of sensory-evoked responses, which are associated with stimulus features and perception, respectively. However, our understanding of how the brain state influences the sensory-evoked activation across the mouse cortex remains limited.

In this study, we utilized isoflurane to modulate the brain state and conducted wide-field calcium imaging of Thy1-GCaMP6f mice to monitor the distributed activation evoked by multi-whisker stimulation. Our findings reveal that the level of anesthesia strongly shapes the spatiotemporal features and the functional connectivity of the sensory-activated network. As anesthesia levels decrease, we observe increasingly complex responses, accompanied by the emergence of the late component within the sensory-evoked response. The persistence of the late component under anesthesia raises new questions regarding the potential existence of perception during unconscious states.

## INTRODUCTION

Sensory information needs to be broadcasted among distributed brain areas to be integrated and used to control behavior^1–3^. Since the pioneering work from the Petersen’s group, showing the local propagation of the activity from a single barrel to the entire barrel field^4^, there has been a growing interest in the integration of sensory information beyond the primary responsive area. Indeed, distributed aspects of cortical information processing over the entire dorsal cortex were identified as a central mechanism for perceiving peripheral stimuli. One of the first studies to demonstrate the importance of integration was by Guo and colleagues, who established the necessity of serial information flow from sensory to motor areas for perceptual decision-making^2^. The propagation from sensory regions to the entire dorsal cortex was then explored by several groups using wide-field imaging^1,5–8^. Notably, a strong dependence of the evoked spatiotemporal dynamics on the brain state was demonstrated by comparing wakefulness and anesthesia^9–12^. This is in line with a significant body of evidence positing that anesthesia is primarily linked to the breakdown of long-range projections between cortical areas, leading to the impairment of cortical information integration^13,14^. According to the integrated information theory (IIT), disruption of this large-scale integration process is linked to a reduction in the complexity of the locally-evoked cortical response^15^. Nevertheless, studies of the complexity associated with a peripheral stimulus, and how this is modulated by the brain state, are largely missing.

It has been demonstrated that in response to a sensory stimulus, the cortex exhibits an early transient response followed by a late response in both awake and anesthetized states ^1,16–19^. While the early component of the sensory response reliably represents the stimulus identity, the late component correlates with the stimulus perception and is independent of stimulus modalities^16,17^. Indeed, blocking the late component by optogenetic inhibition in the primary sensory area impairs the perceptual report^18^. Later studies demonstrated that both the early and late responses are broadcasted across the cortex^1^. Specifically, the spatiotemporal pattern of the late response is different from the early response and can be modulated by experience^16^. Considering the link between delayed responses and sensory perception, one might expect that the former would be diminished in unconscious states, such as during anesthesia. In fact, studies have shown that anesthesia significantly influences responses in the primary sensory area, indicating that both early and late components are affected^20^. However, there is a lack of understanding on how the early and late responses are modulated by the level of anesthesia across the entire cortex and if there is a threshold for the occurrence of the late response.

In this study, we investigated the brain-state dependence of the large-scale dynamics engaged by multi-whisker stimulation using wide-field calcium imaging of Thy1-GCaMP6f mice. Initially, we examined the connectivity structures within the sensory-activated cortical network across four different brain states: wakefulness and three distinct levels of anesthesia. Then, we delved into how the brain state affects the complexity of the sensory-evoked response, and employed a mean-field model to investigate potential mechanisms underlying the experimental findings. Finally, we mapped the spatiotemporal features of the distributed sensory response, both from average and from single trial activity, allowing us to address the long standing question of whether the late response remains observable irrespective of the anesthesia level.

## RESULTS

This study aimed to evaluate how the brain state modulates the spatiotemporal integration of the cortical response to sensory stimulation. We investigated sensory-evoked cortical activity in four separate brain states - three anesthesia levels and the awake state. To this end, we performed widefield calcium imaging of the entire dorsal cortex in transgenic mice expressing GCaMP6f in excitatory neurons (RRID:IMSR_JAX:025393) (Fig. 1a). The subjects were either awake or under three levels of anesthesia: deep, medium, and light. The isoflurane concentration was adjusted within the range of 1.3% to 1.7% to achieve the desired levels of anesthesia which were cross-validated based on the features of the up and down states extracted from the spontaneous calcium activity traces.

**Figure 1.**
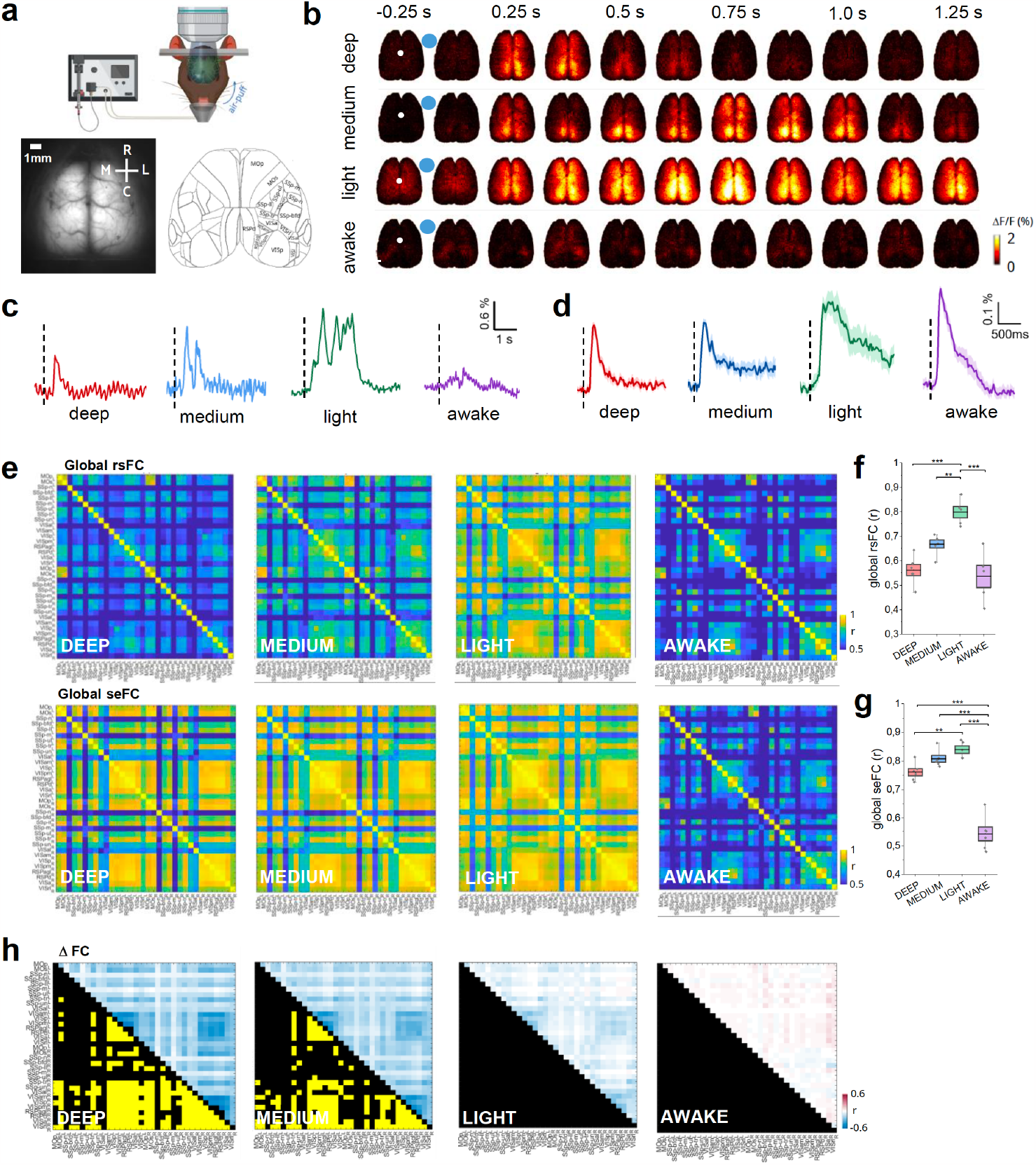
Brain state dependence of the sensory-evoked functional networks. **(a)** Schematic representation of the whiskers stimulation experiment (upper panel). Field of view of the wide-field microscope (lower panel, left) and Allen Mouse Brain atlas parcellation (lower panel, right). **(b)** Representative image sequences showing the cortical activity in different brain states. White dots represent bregma and blue dots represent the stimulus time. **(c)** Single-trial time series showing the cortical response to whisker stimulation in different brain states. The traces represent the average activity of all the pixels in the masked field of view. Vertical dashed lines represent the stimulus time. **(d)** Average traces showing the sensory-evoked activity in the primary sensory area, SSp-bfd, in awake mice and under increasing levels of isoflurane. **(e)** In the upper plan Correlation matrices (Pearson’s correlation, Fisher’s z-transformed) showing the rsFC in the four brain states from deep anesthesia (left) to wakefulness (right). In the lower plan sensory-evoked functional connectivity matrices **(f)** Box plots displaying the global rsFC calculated averaging the correlation values of all paired cortical regions. **(g)** Box plots displaying the global seFC calculated by averaging the correlation values of all paired cortical regions. One-way ANOVA with Bonferroni correction **(h)** Matrices displaying differences between rsFC and seFC for all the four brain states. The quantification of the differences is shown in the upper part of the matrices and the corresponding statistical significance is reported in the bottom part (black square = non-significant; yellow square = Benjamini– Hochberg corrected P < 0.05). N mice = 6

The calcium imaging recordings revealed a distributed and asynchronous activation in the awake state and a synchronous and global activation in the light state, which became progressively more localized in medium and deep anesthesia (Suppl. Fig. 1a). Consistent with previous studies^12,21^, all anesthetized states exhibited bistable activity (Suppl. Fig. 1b) with shorter down states and longer up states as the anesthesia decreased (Suppl. Fig. 1c-d). Interestingly, the duration of the up-states in light anesthesia is similar to those reported in micro-arousals^22^. The upstate frequency was comparable in light and medium states but dramatically decreased in deep anesthesia (Suppl. Fig. 1e).

### Brain state-dependent formation of cortical functional clusters in response to sensory stimulation

Unilateral multi-whisker stimulation elicited a local and transitory response in awake subjects, which was replaced by a persistent and global response across the whole cortex when mice were exposed to a low dose of isoflurane anesthesia (Fig. 1b,c). Peripheral stimulation in medium and deep anesthesia resulted in a progressively shorter and more localized activation, primarily engaging the medial associative regions (Fig. 1b). At the level of the primary sensory area (SSp-bfd), the averaged time series show a more stable activation profile across brain states (Fig. 1d). The spatiotemporal patterns evoked by sensory stimulation, although time-locked to the air puff, presented similarities with the resting state activity under anesthesia. We sought to investigate if the functional connectivity of spontaneous and evoked activity shared similar clusters, and if they were brain-state dependent. Previous studies have reported differences in resting-state functional connectivity (rs-FC) between awake and anesthetized states induced by various anesthetic agents^23^, but the correlation structures elicited by sensory stimulation, or sensory-evoked Functional Connectivity (seFC), are rarely addressed. Within this framework, we examined the brain state dependence of the functional connectivity of excitatory neurons elicited by sensory stimulation and compared it to the resting state. Pearson’s correlation (Fisher’s z-transformed) was computed between all paired cortical regions and expressed as a matrix for both resting-state (rs) and sensory-evoked (se) conditions per brain state (Fig. 1e). Consistent with previous studies ^24^, rsFC increased with decreasing anesthesia levels for all cortical pairs, which was reflected in the global rsFC (Fig. 1f). Transitioning to the awake state resulted in a drastic decrease in rsFC to levels comparable to deep anesthesia (Fig. 1f). Conversely, whisker stimulation increased the seFC of deeper anesthesia levels (Fig. 1g). Accordingly, the difference matrices highlighted the emergence of a cluster of regions significantly different from the resting state in deep and medium anesthesia. This functional cluster was primarily associated with an increased coactivation of both intra- and inter-hemispheric seFC of posterior regions (Fig. 1h). Interestingly, this increase in FC was lost under light anesthesia and in awake mice. These results indicate that the connectivity of cortical regions involved in processing incoming sensory information (seFC) significantly diverges from spontaneous activity (rsFC) only under deeper anesthesia. seFC in wakefulness is substantially different from all anesthesia levels.

### The complexity of evoked responses decreases as the level of anesthesia increases

To determine how anesthesia affects the propagation of information across the brain, we computed the Perturbational Complexity Index (PCI) of the sensory-evoked response from wide-field calcium data and the computational model. The spread of sensory-evoked activity decreases with increasing levels of anesthesia (Fig. 2a), consistent with results in humans25. Using a whole-brain model of the mouse cerebral cortex (Fig. 2b), we simulated different activity states corresponding to asynchronous (wake-like) dynamics (Fig. 2c; top trace) and different slow-wave oscillatory states (Fig. 2c; bottom traces; Fig. 2d for power spectra), using a similar procedure as in a previous model of human slow waves where the transition is simulated by tuning the spike-frequency adaptation26. The response to cortical stimulation is shown in Fig. 2e, in the same asynchronous and slow-wave states. This response was quantified using PCI, and as found in the experiments, the PCI decreased with the strength of slow waves, showing that the spread of the response is diminished in slow-wave states in both model and experiments (Fig. 2f).

**Figure 2.**
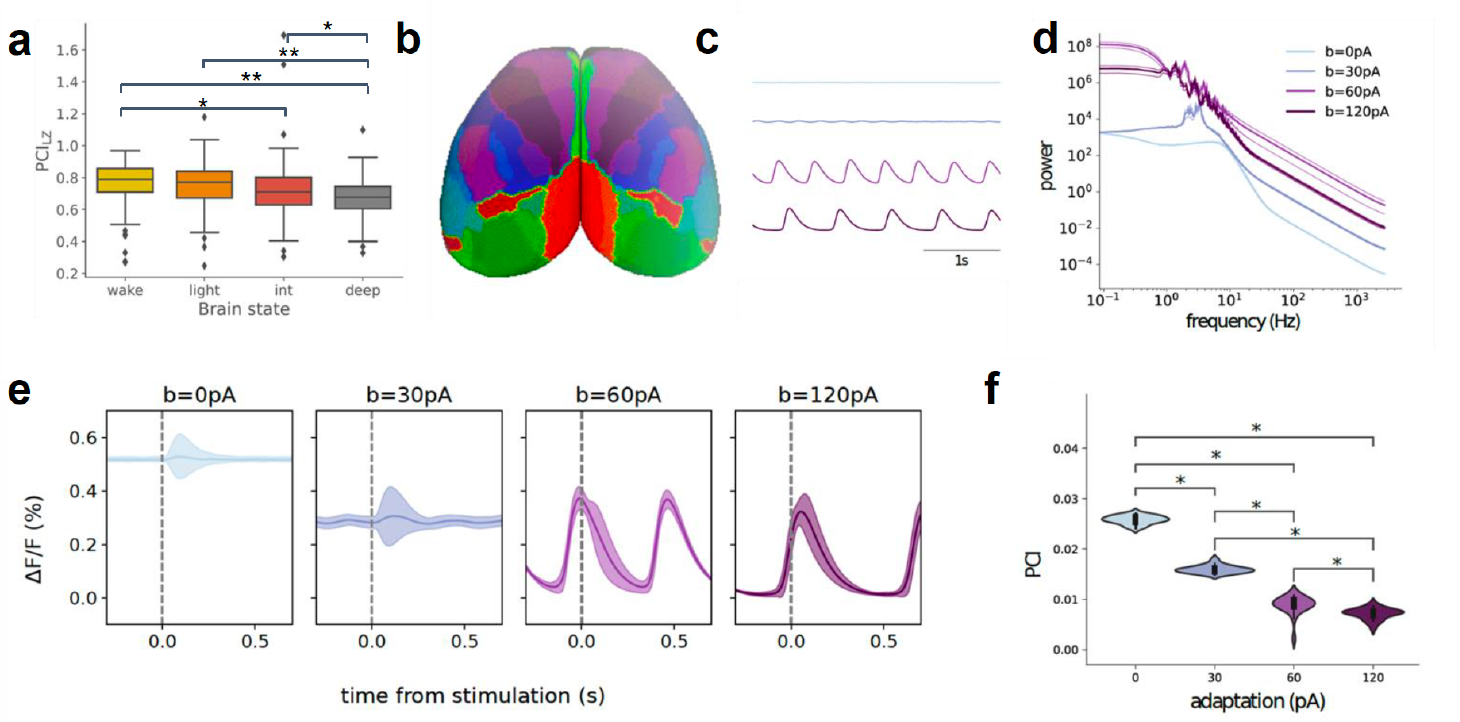
The complexity of the sensory evoked response is brain-state dependent. **(a)** The Perturbational Complexity Index (PCI) of the sensory-evoked response from wide-field calcium data decreases with increasing levels of anesthesia. **(b)** Simulated PCI is calculated from the TVB-Adex model of the entire cortical surface, parcellated according to the Allen Brain mouse atlas. **(c)** Transition from asynchronous (top) to slow-wave activity (bottom), as obtained by augmenting the strength of spike-frequency adaptation in the model (0, 30, 60 and 120 pA from top to bottom). **(d)** Corresponding power spectra showing the shift towards low frequencies. **(e)** Responses to cortical stimulation in different states described in **(c). (f)** PCI calculated from the model for increasing levels of adaptation.

### The spatiotemporal features of the sensory information broadcasting among cortical regions depend on the brain state

Our results demonstrated a high similarity in seFC features across all anesthesia levels. However, the complexity (PCI) of the stimulated activity was significantly brain-state dependent, indicating increasing levels of differentiation and integration with decreasing levels of anesthesia. Therefore, we sought to investigate these features in the spatiotemporal propagation of the evoked response. Consistent with previous studies^1,6^, monolateral whisker stimulation evoked a distributed cortical response with brain state dependency. We analyzed seventeen representative cortical regions per hemisphere to reconstruct topographic response maps based on the mean response. The temporal profile of the evoked calcium transients in the barrel cortex contralateral to the stimulus (ss-bfd_R_) exhibited distinct characteristics for each of the four brain states analyzed (table 1). Specifically, ss-bfd_R_ showed consistent onset, rising slope, and maximum amplitude across all brain states, while the time to peak was longer in light anesthesia and the response duration was longer for medium and light anesthesia compared to the other brain states (table 1-4).

The integration of the response throughout the other cortical regions varied significantly depending on the brain state. While the onset latency was uniform across all brain states (table 1, Fig. 3a), the rise slope of the majority of investigated areas was significantly higher in deep anesthesia compared to wakefulness and gradually decreased moving to medium and light anesthesia (table 2, Fig. 3b). Conversely, the time to peak was significantly longer only in light anesthesia for most cortical areas examined (table 3, Fig. 3c). Notably, the time to reach the peak displayed substantial regional variance in light anesthesia and wakefulness, whereas it was relatively consistent across brain areas in deep and medium anesthesia (table 3, Fig. 3c). Moreover, compared to wakefulness, the three anesthetized states exhibited higher amplitude of the response peak per cortical area (table 4, Fig. 3d). Finally, deep and awake states displayed shorter response duration compared to light and medium anesthesia, which were characterized by a more persistent response (table 5, Fig. 3e).

**Figure 3.**
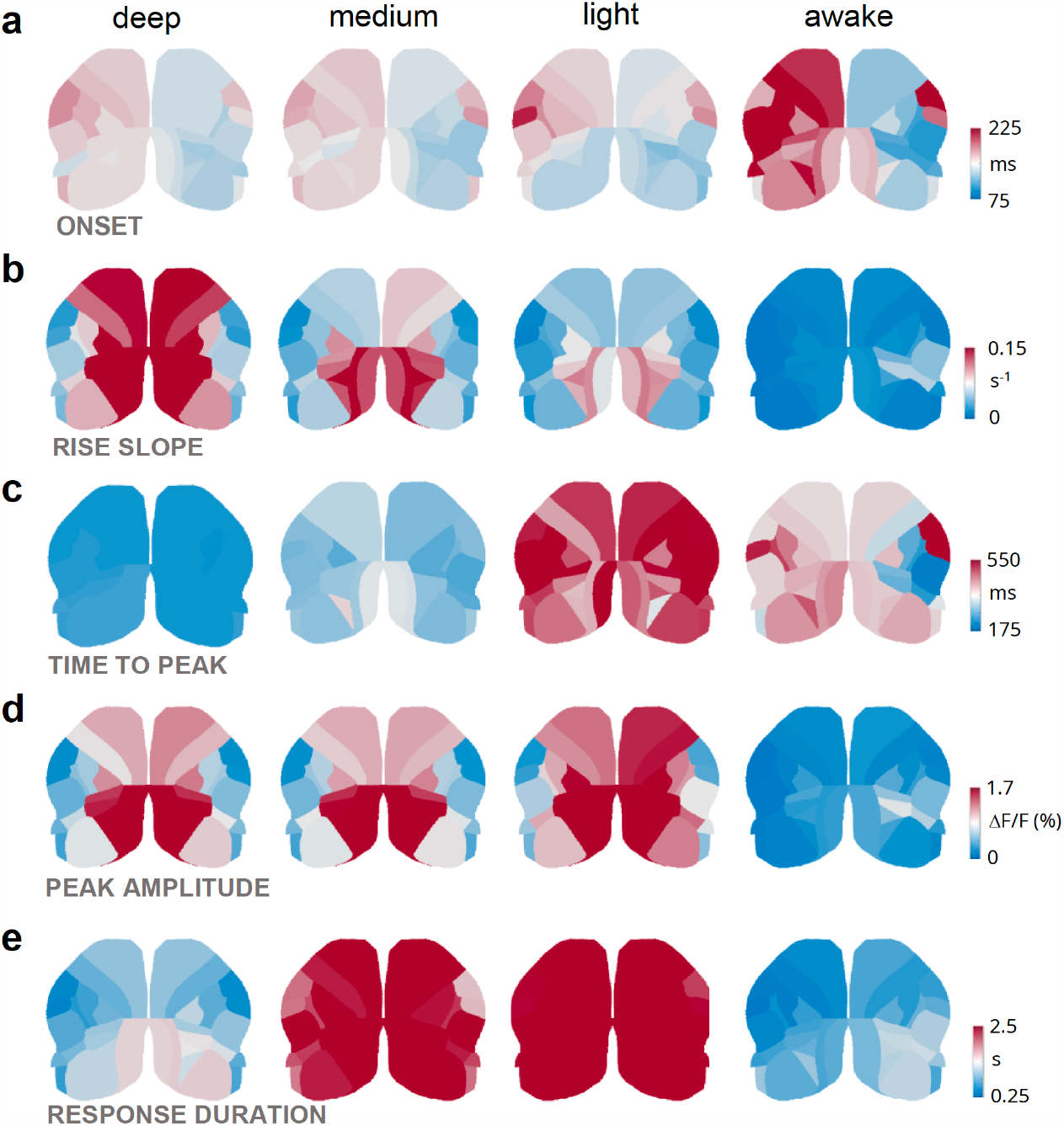
Brain state dependence of the spatiotemporal features of the distributed sensory-evoked response. **(a-e)** Cortical maps registered to the Allen Mouse Brain Atlas displaying the quantification of the **(a)** onset, (**b)** rise slope, **(c)** time to peak, **(d)** peak amplitude and **(e)** response duration, of the activity in different cortical regions. N mice = 6.

These findings highlight the stability of the average response in the primary sensory area ss-bfd_R,_ suggesting that the cortical representation of stimulus features is stable regardless of the brain state. In contrast, the integration of sensory input across the entire dorsal cortex is strongly influenced by the brain state.

### Slight adjustments to the anesthesia level result in the appearance of the late response to the stimulus

Our results revealed that sensory stimulation engaged an FC cluster that did not differ between medium and deep anesthesia. This similarity was also observed for the activation in the primary sensory area ss-bfd_R_ which we found comparable in these two brain states. However, significant brain state-dependent differences were observed in the complexity of the evoked response and in the integration of the response across the entire cortex. One key feature that distinguished deep and medium was the average response duration, which could be linked to an increased likelihood of the emergence of a late component. Therefore, we hypothesized that the late component could be responsible for the increased complexity and duration of the evoked response in medium anesthesia. Specifically, we wondered if the brain state could selectively modulate different aspects of these two time domains.

To test these ideas, we conducted a single trial analysis of the sensory-evoked response in the same animals under deep and medium anesthesia. We analyzed six representative regions per hemisphere: the primary and secondary motor cortices (M1 and M2), primary somatosensory area of the barrel field (SS-bfd), primary somatosensory area of the trunk (SS-tr), primary visual cortex (V1) and retrosplenial cortex (RSP) (Fig. 4a). The exemplary traces showed that a stereotyped early response was present in both anesthesia levels, followed by a late response that was predominantly observed in medium anesthesia and largely absent in deep anesthesia (Fig. 4b). The analysis of the early response revealed that the onset was symmetrical between hemispheres and was not dependent on the anesthesia level (Fig. 4c). Interestingly, the time to peak and full width at half maximum (FWHM) were increased in medium compared to deep anesthesia, with a prominent contribution from the contralateral hemisphere to the stimulus (Fig. 4d,e). Although the peak amplitude of the early component exhibited asymmetry within a single brain state due to higher values in the contralateral hemisphere to the stimulus, it did not significantly differ between the two anesthetized levels indicating that it was not sensitive to the brain state (Fig. 4f). Moreover, both the rise and decay slopes were consistently lower on the contralateral hemisphere in most of the analyzed regions and the rising slope also showed higher values in deep compared to medium anesthesia (Fig. 4g,h). Overall, these features indicate that sensory stimulation leads to a slower and longer early response in medium anesthesia while maintaining the same amplitude as in deep anesthesia.

**Figure 4.**
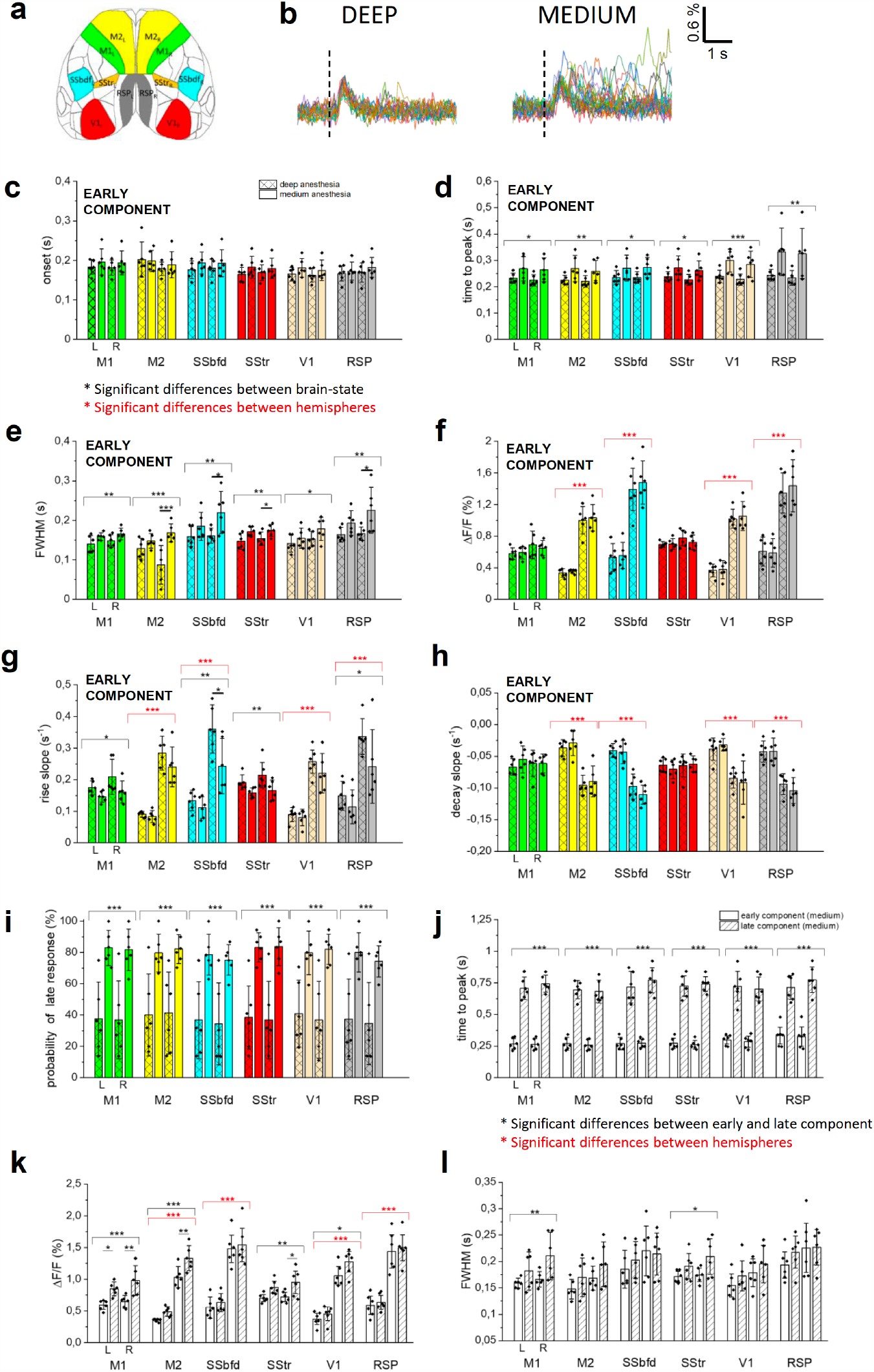
Small variations in the anesthesia level determine substantial changes in the late component of the sensory-evoked response. **(a)** Allen Mouse Brain atlas parcellation used to identify the cortical regions for single-trial analysis (M1 – primary motor cortex; M2 – Secondary motor cortex; SSbdf – primary somatosensory barrel field cortex; SStr - primary somatosensory cortex trunk; RSP – Retrosplenial cortex; V1 – primary visual cortex; L – left; R – right). **(b)** Sensory-evoked calcium activity in the Barrel field cortex under deep (on the right) and medium (on the left) anesthesia in the same representative animal (dashed line indicates the timing of the stimulus, n stimuli=11). Column chart showing onset time **(c)**, time to peak **(d)**, FWHM **(e)**, ΔF/F% **(f)**, rise slope **(g)** and decay slope **(h)** of the early component for six cortical regions in the left (L) and in the right (R) hemisphere under deep (patterned columns) and medium (empty columns) anesthesia. Whisker length is the standard error. **(i)** Box plot showing the probability of late response for six cortical regions in the left (L) and in the right (R) hemisphere under (patterned columns) and medium (empty columns) anesthesia. Whisker length is the standard error. Box plot showing comparing time to peak **(j)**, ΔF/F% **(k)** and FWHM **(l)** of the early (empty column) and late component (patterned column) for six cortical regions in the left (L) and right (R) hemisphere under medium anesthesia. N mice = 6. Two-way ANOVA with Bonferroni correction (* p < 0.05; ** p < 0.01; *** p < 0.001)

We previously highlighted the presence of a late component, primarily observed in the medium state. The probability of evoking a late response dramatically decreased from medium to deep anesthesia (Fig. 4i). Consequently, in our further analysis, we focused on the comparison between the early and late components within medium anesthesia. The late response reached the maximum peak almost uniformly across brain regions and hemispheres, approximately 500 ms after the early response (Fig. 4j, Suppl. Fig. 2a). However, the amplitude of the late response exhibited a precise topographic modulation, with higher values in the contralateral hemisphere, except for BF and RSP (Fig. 4k, Suppl. Fig. 2b). The duration of the early and late activations, measured by FWHM, was mostly comparable across all cortical regions, with a few exceptions (M1 and SSTr) (Fig. 4l).

Considering that the late response has been previously linked to the subjective perceptual report of the stimulus^18^, these findings might imply that medium anesthesia, as opposed to deep anesthesia, is associated with heightened integration of the sensory stimulus. Consequently, even minor adjustments to anesthesia levels results in substantial variations in the distributed cortical processing of the stimulus.

In conclusion, our observations on the late response suggest that the mechanisms underlying stimulus perception may be activated even under medium levels of anesthesia.

## Discussion

Different states of the brain are characterized by alterations in the spatiotemporal pattern of neuronal activity in many brain areas^11^ which can influence responses to incoming sensory information^9,10^. To investigate the interplay of local and distributed cortical responses to sensory stimulation and its dependence on the brain state, we took advantage of wide-field optical imaging of excitatory neurons in Thy1-GCaMP6f mice. We examined the distributed cortical activity evoked by unilateral whisker stimulation across four distinct brain states, including three levels of isoflurane anesthesia and wakefulness.

Our results indicate that increasing levels of anesthesia are associated with (i) the emergence of a cluster of posterior regions exhibiting higher intra- and inter-hemispheric correlation during sensory stimulation (ii) a progressive loss of cortical response complexity (iii) a reduction in the duration of the mean response across all cortical regions, which is arguably due to (iv) the disappearance of a late component of the sensory-evoked response.

Cortical connectivity may reflect distinct patterns of activation in the evoked activity compared to the resting state and could be modulated differently by experience ^27,28^. The vast majority of studies comparing resting state and task-evoked FC are based on human fMRI measures ^29–36^. They show that the rs- and se-FC can share common patterns, being them generated by the activation of partially overlapping networks. Here we compared the seFC to the rsFC and found significant differences only in deeper anesthesia levels. Our findings, which support the concept that the functional connectivity of sensory-evoked cortical responses can significantly diverge from that of spontaneous activity, provide a crucial insight by indicating that this distinction is significantly influenced by the brain state. Surprisingly, the difference between rsFC and seFC becomes more pronounced in deeper anesthesia states, where one would expect more stereotypical activity regardless of the source of activation (internal vs. peripheral). Conversely, the similarity in the coactivation of the cortical network in the wakefulness state indicates that the main driver is likely the constantly changing patterns of activation, regardless of the origin of the signal. Future research is needed to further investigate how sensory experience can modify seFC and how this process is dependent on the level of anesthesia.

The quantification of the spread of information in the different brain states was done using PCI, which was found to globally decrease with the depth of anesthesia (Fig. 2a). This global decrease could be replicated by a mouse whole-brain model, where the PCI was inversely proportional to the level of adaptation (Fig. 2f). This result must be put in perspective with findings in humans, where the evoked activity is maximally complex in the awake state^37^, and the PCI drops to low values during sleep or anesthesia^25^. This result was also replicated by a whole-brain model^26,38^. It must be noted, however, that the present results are different because a sensory input was used, while the model used direct cortical stimulation. This may explain why the PCI variations are larger in the model compared to the experiments, a point that should be checked when more precise models are available to simulate sensory inputs.

In studies investigating the neuronal correlates of sensory stimuli, averaging signals across trials is a common approach to enhance the signal-to-noise ratio and reveal the underlying neural response. However, this averaging approach may mask meaningful time-varying neural fluctuations that can be important for understanding brain function^3,39^. Furthermore, it has been demonstrated that the trial-to-trial variability of sensory-evoked responses is dependent on the cortical state^9,40,41^. To overcome these limitations, we utilized the high signal-to-noise ratio and fast dynamics of the GCaMP6f indicator in combination with the temporal resolution of our optical setup to explore the sensory-evoked dynamics at the single-trial level. This allowed us to characterize the brain-state dependence of the late response to sensory stimulation and uncover specific features of the early and late components that are hard to disentangle from the average response.

The impact of brain state on the coding of a stimulus has mainly been investigated at the local level, particularly in the primary sensory area ^9,16^. These studies primarily focus on the temporal organization of locally evoked activity during anesthesia and analyze complexity, onset, and duration, with findings comparable to our observations for the medium anesthesia level ^16^.

In the primary sensory area (SSbfd contralateral to the stimulus), our results demonstrate that the average response is stable in terms of amplitude, onset, and rise time across all brain states. Nonetheless, in other cortical regions, most of the spatiotemporal features are markedly influenced by the brain state. This suggests that it is predominantly the late processing and its broadcasting throughout the dorsal cortex that are dependent on the brain state, as demonstrated by the significant appearance of a late response only beyond a certain threshold of isoflurane level.

These findings align with previous work suggesting that the organization of the early-late component in sensory-evoked activity can be modulated by the brain state ^16,42^. We observed that the early component of the evoked response exhibits a longer time to peak and duration (FWHM) across all cortical regions in medium anesthesia compared to deep anesthesia (Fig. 4d, e). Additionally, the early component shows strong asymmetry in M2, SSbfd, and RSP but similarities between brain states in terms of amplitude (Fig. 4f), indicating differences in the perception of stimulus features between hemispheres but no influence of the anesthesia level on amplitude. In contrast, the late component is strongly brain-state dependent, as the probability of evoking a late response dramatically drops in deep anesthesia (Fig. 4i). Previous studies have demonstrated that the late activity phase is critical for perception, as silencing or blocking it interferes with stimulus perception or detection ^17,18,43^. Therefore, our results suggest that the perception of the stimulus could still be present under anesthesia and can disappear only at very deep anesthesia levels.

It has been suggested that stimulus perception requires the recirculation of activity through recurrent local and horizontal connections, which typically influences the late phase of the response^17^. Recent studies have indicated that neurons in higher areas are responsible for sending cortical feedback, which is reflected in the late response in the primary sensory area^44–46^. Our findings show that the late response exhibits a greater delay in SSbfd and RSP compared to M2 in the contralateral hemisphere to the stimulus, suggesting an anteroposterior sequence of activation starting from neurons in the secondary motor cortex and later reaching the primary sensory region. The high amplitude of the late component in SSbfd and RSP, followed by M2 and V1, indicates their substantial involvement in cortico-cortical feedback. Given that the late response’s amplitude and activity recirculation are both related to stimulus perception, our results support the hypothesis that stimulus perception is a distributed process involving multiple cortical areas beyond the directly stimulated one (Suppl. Fig. 2 and Fig 4), with specific involvement of M2, SSbfd, and RSP in the late response.

In summary, our findings demonstrate that the majority of features of sensory-evoked activity that are sensitive to the brain state belong to the late component of the cortical response. The perception of the stimulus is strongly influenced by the complexity of the brain state and likely depends not only on the local manifestation of a late response but also on how this response is propagated and integrated by other cortical regions.

## Material and Methods

### 1. Animals

All experiments were performed in accordance with the guidelines of the Italian Minister of Health (aut. n. 857/2021). A total of 9 mice (6-12 months), C57BL/6J-Tg(Thy1-GCaMP6s)GP4.12Dkim/J^47^ heterozygous of both sexes were used.

### 2. Intact-Skull preparation

Mice were anesthetized with isoflurane (3% for induction, 1–2% for maintenance) and placed in a stereotaxic apparatus (KOPF, model 1900). Ophthalmic gel (Lacrilube) was applied to prevent eye drying, body temperature was maintained at 36°C using a heating pad and lidocaine 2% was used as local anaesthetic. The skin and the periosteum were cleaned and removed. Bregma was signed with a black fine-tip pen. A custom-made aluminium head-bar placed behind lambda and the exposed skull were sealed using transparent dental cement (Super Bond C&B – Sun Medical). After the surgery, mice were recovered in a temperature- and humidity-controlled room, with food and water ad libitum for two weeks before recordings.

### 3. Imaging setup

The setup was slightly modified from ^48,49^. Wide-field imaging was performed through the intact skull using a custom-made microscope. The excitation source for GCaMP6f was a blue-light beam of emitting diode (470nm LED light, M470L3 Thorlabs, New Jersey, United State). The excitation band was selected by a bandpass filter (482/18 Semrock, Rochester, New York, NY, USA). The light beam was deflected by a dichroic mirror (DC FF 495-DI02 Semrock, Rochester, New York, NY, USA) on the objective (TL2X-SAP 2 Super Apochromatic Microscope Objective, 0.1 NA, 56.3 mm WD).

Reflectance images were acquired using a light source positioned at 45° incident to the brain surface (530 nm LED light, M530L4; Thorlabs, New Jersey, United State). The excitation band was selected by a bandpass filter (525/50 nm filter). Both the fluorescence signal and the reflectance changes were collected through a band-pass filter (525/50, Semrock, Rochester, New York, USA) and collected by a high-speed complementary metal-oxide semiconductor (CMOS) camera (ORCA-Flash4.0 V3 Digital CMOS camera / C13440-20CU, Hamamatsu). Reflectance data are acquired interlaced with fluorescence data by using a camera configured to acquire images in synchrony with two strobing LEDs (sampling rate: 40Hz per LED).

### 4. Awake imaging

After the post-surgical recovery period (3 days), mice (n = 5) were acclimatized to the head-fixation for two consecutive days (15 min a day/mouse). This was followed by 50-60 minutes of imaging session in which both spontaneous (38 s-long, 5 repetitions) and perturbed activity (24 s-long, 30 repetitions) were recorded.

### 5. Anesthetized imaging

Mice were anesthetized by isoflurane to investigate three brain states classified according to the isoflurane concentrations in DEEP anesthesia (1.71 ± 0.08 %, n = 6), MEDIUM anesthesia (1.49 ± 0.06 %, n = 6) and LIGHT anesthesia (1.32 ± 0.12 %, n = 5). Each anesthesia level was maintained for 60 minutes, and recordings were consistently monitored to conserve a stable slow-oscillatory regime. Spontaneous activity recordings (38 s-long, 5 repetitions) and perturbed activity recordings (24 s-long, 30 repetitions) were acquired in the same daily session per brain state. Deep and medium anesthesia were recorded consecutively on the same imaging session per mouse, starting from the higher isoflurane concentration to the lower. Light anesthesia was instead recorded on a different day. During the whole anesthetic treatment, body temperature was maintained at 37°C by a feedback-controlled thermostatic heating pad.

### 6. Whisker stimulation

Stimulation was delivered to the left whiskers through a tubing system using an electrically gated pressure injector (Picospritzer III—Science Products). Whiskers were deflected ∼1 cm in the rostrocaudal direction. A single stimulation trial consisted of 24 s designed as 13 s rest, 1 s of stimulation with a blowing time of 60 ms, and 10 s rest. Each experiment session consisted of 30 trials per animal (n = 9 mice).

### 7. Data analysis

Images were recorded at a frame rate of 40Hz per LED, with a resolution of 512 x 512 pixels and a FOV of ∼ 12 x 12 mm (depth 16-bit) via custom-made software. All data analyses were performed in MATLAB (MathWorks), Phyton, ImageJ, and Origin.

ΔF/F was computed for each pixel, where ΔF was the intensity value of that pixel in a specific time point and F was the mean fluorescence intensity of the signal across time for the spontaneous activity or the fluorescence intensity of the pre-stimulus signal for perturbed activity. Then, hemodynamic correction was performed as described by ^50^. Briefly, using the ratiometric approach:

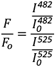

Where F/F_0_ is the final corrected GCaMP6 time series for a given pixel, I^482^ refers to the detected fluorescence signal, I^525^ is the reflectance signal.

To dissect the contribution of each cortical area for both awake and anesthetized activity, the plane of imaging was projected to the surface of the Allen Institute Mouse Brain Atlas (www.brain-map.org) as described by ^49^. A mask was applied to exclude areas laying on the most lateral parts of the mouse cortex and the fluorescence time course was measured by averaging all pixels within individual areas. The neocortex parcellation created 17 areas for each hemisphere, for a total of 34 areas. The abbreviations and extended names for each area are as follows: MOs, secondary motor area; MOp, primary motor area; SSp-bfd, primary somatosensory area, barrel field; SSp-ll, primary somatosensory area, lower limb; SSp-m, primary somatosensory area, mouth; SSp-n, primary somatosensory area, nose; SSp-tr, primary somatosensory area, trunk; SSp-ul, primary somatosensory area, upper limb; SSp-un, primary somatosensory area, unassigned; RSPagl, retrosplenial area, lateral agranular part; RSPd, retrosplenial area, dorsal part; VISrl, rostrolateral visual area; VISa, anterior visual area; VISal, anterolateral visual area; VISam, anteromedial visual area; VISp, primary visual area; and VISpm, posteromedial visual area. The suffixes L and R were added to refer to the left (ipsilateral with respect to the whisker stimulation) or right hemisphere (contralateral with respect to the whisker stimulation) (e.g.., MOp_L, MOp_R).

To quantify the durations of the up states, the calcium signal had to exceed a critical threshold defined by 2 SD the lowest mean activity computed across the entire cortex. Then, the time between two up states was used as down state duration. To avoid random oscillation detection, both the up and down state durations were set to a minimum of 75ms. This threshold was heuristically set after a visual inspection of the signal. Individual perturbed activity trials were discarded prior to analysis if there was an ongoing up state at the time of stimulation in anesthesia.

#### Functional connectivity and network analyses

Three different types of connectivity analyses were performed to assess the impact of the brain state on the FC in both spontaneous and perturbed activity: global, inter- and intra-hemispheric connectivity analysis, and area pair-wise analysis. These analyses were carried out on 2-sec post-stimulus windows for perturbed activity and on 30 randomly selected 2-sec windows for spontaneous activity. Functional connectivity was calculated as the Pearson’s correlation coefficient between signal time series of various cortical regions at the single-trial level. Connectivity scores were averaged over trials within conditions, but independently for each mouse. These scores were subsequently averaged across mice within each brain state. In order to visualize the matrix of differences, the average FC scores of one brain state were subtracted from the corresponding average FC scores of another brain state. To enable parametric statistical testing, correlation coefficients were additionally Fisher z-transformed.

Right and left intra-hemispheric FC were quantified by averaging correlation values within all right and left-hemisphere cortical regions, respectively. Inter-hemispheric FC was measured by averaging correlation values between all right- and left-hemisphere region pairs. By averaging every combination value previously obtained, global FC was calculated. Intra/inter FC ratio was performed by dividing the two average values per mouse. For area pair-wise analysis multiple comparisons were corrected by Benjamini–Hochberg procedure. The significance of the correlation analysis was set at a threshold of (Benjamini–Hochberg) corrected P < 0.05.

No statistical methods were used to predetermine the sample size.

#### Quantitative analysis of perturbed activity

Before processing, the calcium response evoked by stimulation was averaged across 30 trials per cortical area per mouse and per brain state. The onset of the response was defined as the time when the signal crosses a threshold corresponding to the averaged baseline amplitude (1 s before the start of the stimulus) + 2 SD. Only trials with post-stimulus amplitude higher than the threshold in a 2-second window were included in the analysis.

The parameter “time to peak” was quantified as the time interval (s) between the stimulus pulse and the peak response. The response duration was the time interval between the onset of the response and the point at which the activity decreased under the previously determined threshold in a 2-second window. Then, the maximum response was calculated at the time to peak and the FWHM (full width for half-maximal amplitudes) was the interval between time points for half-maximal amplitudes at the ascending and the descending arm of a peak. The rise and decay slope were calculated by a least-squares fit to the trace during the transient.

For only deep and medium anesthesia a single trial analysis was performed to quantify the first and second peak at a single stimulus level and then the analyzed data were averaged across trials per animal. Before processing, single activity traces were temporally smoothed to improve SNR. Six principal areas per hemisphere were selected: MOs, secondary motor area; MOp, primary motor area; SSp-bfd, primary somatosensory area, barrel field; SSp-tr, primary somatosensory area, trunk; VISp, primary visual area; and RSPd, retrosplenial area, dorsal part. The suffixes L and R were added to refer to the left (ipsilateral with respect to the whisker stimulation) or right hemisphere (contralateral with respect to the whisker stimulation) (eg., MOp_L, MOp_R). Only the fraction of trials detected as responsive for the first peak were used to calculate the probability of the second peak.

#### PCI analysis

The Perturbational Complexity Index (PCI) was computed in response to an external stimulus following the method proposed by ^25^. The PCI is the ratio of two quantities: the Lempel-Ziv algorithmic complexity and the source entropy. To compute both quantities, the calcium signal intensity was calculated and binarized to produce significance vectors s(t). Different trials of the same stimulus were aligned to stimulation time, considering the 600 ms before and after stimulus onset. Then, each node over the hemisphere is re-scaled by mean and standard deviation given by pre-stimulus activity, averaged over nodes. After all pre-stimulus signals are randomized across time bins, this procedure is repeated 500 times. The threshold for significance T is then given by the one-tail percentile of the maximum absolute value over all repetitions within a series of 20 trials. For each trial of those 20 trials, we can then write s(t) = 1 whenever post-stimulus calcium signal C(t) > T and S(t) = 0 otherwise.

The Lempel-Ziv complexity LZ(S) is the length of the ‘zipped’ vector S(t), i.e. the number of possible binary ‘words’ that make up the binary vector S(t). Briefly, S(t) is sectioned successively into consecutive words of between one and N_t characters where N_t is the total length of S(t). Scanning sequentially through all words, each newly encountered word is added to a ‘dictionary’, and LZ(S) is the total number of words in the dictionary at the end of the procedure.

The spatial source entropy H(S) is given by:

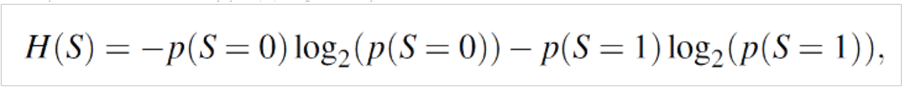

where log_2_ denotes the base-two logarithm. The PCI can then be expressed as

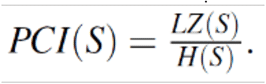

In the model, the PCI was calculated analogously to experimental data, taking as signal the calcium signal generated by the TVB model in each region. Thus, the nodes were given by the different cortical regions of the TVB model.

### 8. Statistical analysis

All statistical analysis was performed in OriginLab (2019). Data are shown as mean ± s.e.m. For multiple comparisons one-way or two-way ANOVA was used and Bonferroni correction was applied for post-hoc t-tests. For the PCI analysis, one-way ANOVA with post-hoc Tukey’s HSD test were used. In the figures, significance levels are represented with the following convention: ∗ for p<0.05; ∗ ∗ for p<0.01, ∗ ∗ ∗ for p<0.001.

## Acknowledgements

This work has been funded by the European Community (Human Brain Project, H2020-785907 and H2020-945539). In addition, this project has been supported by the Italian Ministry for Education in the framework of Eurobioimaging (ESFRI research infrastructure) - Advanced Light Microscopy Italian Node, and from the European Union’s Horizon 2020 Framework Programme for Research and Innovation under grant agreement No. 654148 Laserlab-Europe.

## Author contributions

ALAM, AD, FSP conceived the study; EM and FR did the experiments; EM analyzed experimental data; AS helped in developing the analysis tools; GM developed the software for the imaging setup; NTC participated to the analysis and realized the computational model; FSP contributed funding and resources. EM, FR, NTC, AD, ALAM discussed the results and participated to write the paper; ALAM and AD co-supervised the study.

## Competing interests

The authors declare no competing interests.

## Data Availability

The data in the main figures and tables that support the findings of this study are available from the corresponding author upon reasonable request. Supplementary information is available at communications biology’s website.

## Code availability

The codes used in this study for analyzing calcium imaging data are available from the corresponding author upon reasonable request.

## Notes

### Competing Interest Statement

The authors have declared no competing interest.

