## Supplementary Figures for "Cortical mapping of sensory responses reveals strong brain-state dependence of the late component"

### Mapping the brain-state-dependent sensory response across the mouse cortex

### Supplementary Figures

**Supplementary Figure 1**

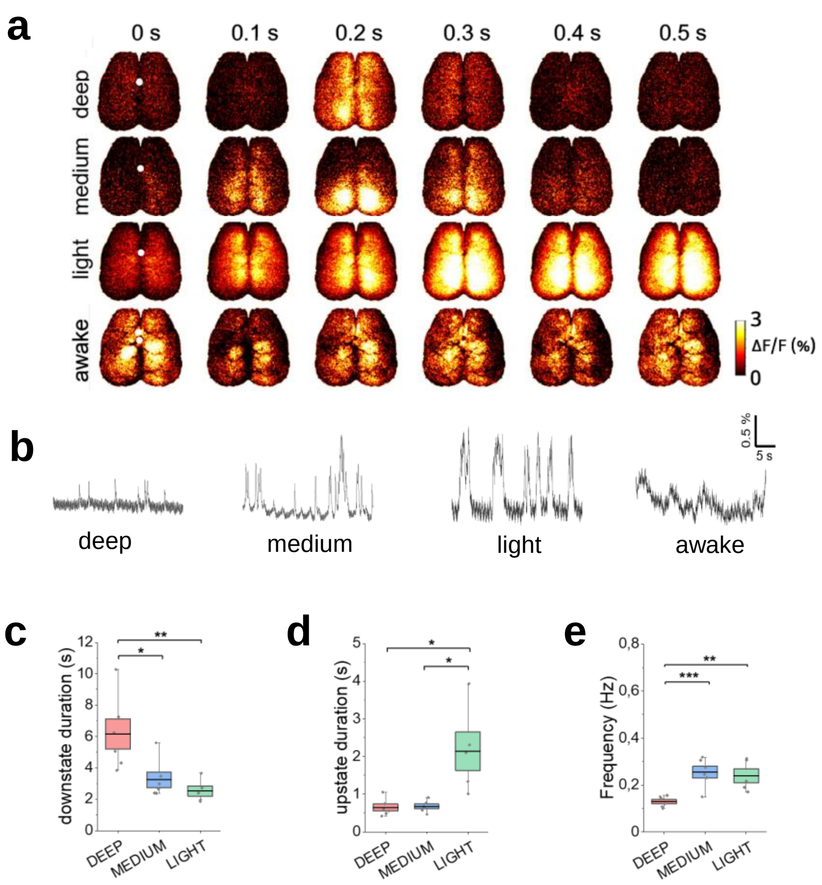


**Supplementary Figure 1. Brain state dependence of the distributed cortical response in resting state conditions.** (a) Representative sequences of resting-state cortical activations in four brain states: awake, light, medium and deep isoflurane anesthesia (11.6 mm x 11.6 mm, frame rate 40 Hz). Scale bar, 1 mm. White dots indicate bregma. (b) Representative calcium traces averaged over the entire cortical surface in the four brain states. (c-e) Characterization of the four brain states in terms of average up states duration (c), down states duration (d), and frequency of up states (e). One-way ANOVA - Bonferroni correction post-hoc t-tests (* $p < 0.05$; ** $p < 0.01$; *** $p < 0.001$)

**Supplementary Figure 2**

**a**

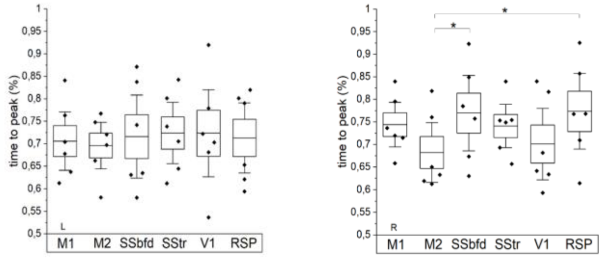


**b**

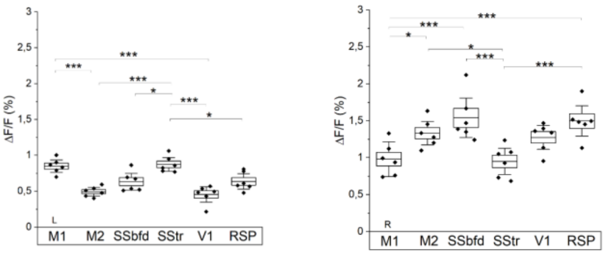


**Supplementary Figure 2**. Box plots displaying the average time to peak (a) and amplitude (ΔF/F) (b) of the late component of the sensory-evoked response for 6 cortical regions on the left hemisphere (L, left panel) and on the right hemisphere in medium anesthesia (R, right panel) (N mice = 6, one-way ANOVA).
